# Dual thermal ecotypes co-exist within a nearly genetically-identical population of the unicellular marine cyanobacterium *Synechococcus*

**DOI:** 10.1101/2020.05.27.119842

**Authors:** Joshua D. Kling, Michael D. Lee, Eric A. Webb, Jordan T. Coelho, Paul Wilburn, Stephanie I. Anderson, Qianqian Zhou, Chunguang Wang, Megan D. Phan, Feixue Fu, Colin T. Kremer, Elena Litchman, Tatiana A. Rynearson, David A. Hutchins

**Author notes:** **Corresponding Author**: David A. Hutchins, Address: 3616 Trousdale Parkway, AHF 207, Los Angeles, CA 90007. **Author Contributions:** Isolates were collected by JDK, PW, SIA, EL, TAR, & DAH, and subsequent culture work and its analysis done by JDK, MDP, EAW, FU, CTK. DNA extraction, sequencing, and analysis performed by JDK, MDL, JTC, QZ, & CW. All authors contributed to the writing.

## Abstract

The extent and ecological significance of intraspecific diversity within marine microbial populations is still poorly understood, and it remains unclear if such strain-level microdiversity will affect fitness and persistence in a rapidly changing ocean environment. In this study, we cultured 11 sympatric strains of the ubiquitous marine picocyanobacterium *Synechococcus* isolated from a Narragansett Bay (Rhode Island, USA) phytoplankton community thermal selection experiment. Despite all 11 isolates being highly similar (with average nucleotide identities of >99.9%, with 98.6-100% of the genome aligning), thermal performance curves revealed selection at warm and cool temperatures had subdivided the initial population into thermotypes with pronounced differences in maximum growth temperatures. Within the fine-scale genetic diversity that did exist within this population, the two divergent thermal ecotypes differed at a locus containing genes for the phycobilisome antenna complex. Our study demonstrates that present-day marine microbial populations can contain microdiversity in the form of cryptic but environmentally-relevant thermotypes that may increase their resilience to future rising temperatures.

**Significance:** Numerous studies exist comparing the responses of distinct taxonomic groups of marine microbes to a warming ocean (interspecific thermal diversity). For example, *Synechococcus*, a nearly globally distributed unicellular marine picocyanobacterium that makes significant contributions to oceanic primary productivity, contains numerous taxonomically distinct lineages with well documented temperature relationships. Little is known though about the diversity of functional responses to temperature within a given population where genetic similarity is high (intraspecific thermal diversity). This study suggests that understanding the extent of this functional intraspecific microdiversity is an essential prerequisite to predicting the resilience of biogeochemically essential microbial groups such as marine *Synechococcus* to a changing climate.

## Introduction

Marine bacteria control most marine biogeochemical cycles (1, 2) and are composed of an estimated 10^10^ species (3). In addition, bacterial species complexes also include numerous strains or ecotypes (4–8). Much of the work documenting the ecological relevance of intraspecific microdiversity has used amplified marker genes such as the 16S rRNA gene, resolved to single base pair differences (9). However, there is still much work to be done to describe the potentially substantial genotypic and (more importantly) phenotypic diversity that exists within groups that are highly similar or even identical at the 16S rRNA level. At higher taxonomic levels, microbial interspecific diversity has a recognized role increasing the stability of biogeochemical cycling and resilience to a changing environment (10). In order to understand the ability of microbial populations to persist and maintain their functional roles under changing thermal regimes, however, it is also important to understand how much unrecognized microdiversity relevant to future warmer temperatures currently exists within microbial populations (11–13).

Efforts to understand the interactions of microbes with the marine environment have often relied on approaches that underestimate or mask intraspecific diversity. For instance, culture-based methods are often limited to a handful of strains that are amenable to cultivation or are currently available in culture collections. These are then used to make generalizations about the activity of a broader taxonomic group (14–16). Sequencing approaches avoid this culturing bottleneck but lack the ability to provide rate measurements. Furthermore, metagenomic or metatranscriptomic analysis pipelines often are unable to discern sequencing errors from rare genotypes or strains (17). In addition, purely sequence-based *in situ* approaches are also limited in the amount of ecotype microdiversity they can reveal, simply because detection relies on observed correlations between relative abundance and ambient environmental parameters (e.g. temperature, nutrients, light). Thus, rare ecotypes with optimal niches that lie outside of current conditions will remain cryptic unless the environment changes. For example, most marine microbial communities will undergo future selection by temperatures exceeding those that they currently experience (18, 19), as current climate models predict that anthropogenic carbon emissions will raise sea surface temperatures ~4°C by the year 2100 (20).

Marine unicellular picocyanobacteria are particularly important to our understanding of how microbially mediated biogeochemical cycling will change with rising sea surface temperatures (SST). The unicellular marine cyanobacterium *Synechococcus* is a major microbial functional and taxonomic group that is found from the equator to high polar latitudes (8, 21). This widespread and diverse genus is responsible for an estimated 16.7% of marine primary production, and is expected to increase in both abundance and distribution as result of climate warming (22). It has also been strongly correlated with carbon export to the deep ocean (23), making this genus an important component of the marine carbon cycle.

In this study, we used multiple temperature incubations of a natural coastal assemblage to enrich for ‘thermal specialist’ strains of *Synechococcus*. We then isolated multiple sympatric strains from the contrasting temperature incubations and characterized their thermal niches, allowing us to recover two co-occuring but distinct thermal phenotypes from a single initial water sample. Finally, we used high-coverage, short read sequencing to obtain high-quality draft genomes for all of the isolates. Upon comparing their assemblies, the two sets of thermally-distinct isolates are nearly identical, with an average nucleotide identity (ANI) > 99.9% demonstrating that these thermotypes belong to the same population. We additionally detected genetic variation that differentiates both thermotypes in a locus coding for the photosynthetic accessory pigment C-phycocyanin. This trait’s relevance is not immediately apparent within the current study’s context, but is nevertheless strongly correlated with the identifiable thermal specializations observed in this population.

## Results

Out of the 11 strains of *Synechococcus* isolated in this study, one originated from our initial cell sorting of the collected seawater (at 22°C), before nutrients were added (Table S1). The other ten isolates were collected from the enrichment experiments, five from an 18 °C enrichments and five from 30 °C. Because the 22°C *in situ* conditions and 18°C experimental treatment represent temperatures that currently occur in Narragansett Bay, these isolates are considered together and are referred to as “cool temperature isolates” and compared against “warm temperature isolates” collected from 30 °C (a temperature exceeding those currently recorded at this sampling site). Despite these considerable differences in temperature, all of the cool and warm temperature isolates shared virtually identical morphologies (Table S1). Picocyanobacteria were also detected in 22 and 26 °C temperature incubation treatments, but cell sorting did not produce any culturable isolates from these incubations.

After growing each of the 11 isolates across multiple temperatures, we generated thermal performance curves for each isolate, with an average R^2^ of 0.81 (±0.14 SD, Figure 1A & S1; Table S2). The average thermal maximum (Tmax) was highest for warm temperature isolates (35.6 °C, ±0.5 SD) compared to cool temperature isolates (33.5 °C, ±0.9; t-test, *p* = 0.005; Figure 1B; Table 1). The optimal growth temperature (Topt) was also higher for isolates from warm temperatures, with a mean of 29.8° (±1.8 SD), than for cool temperature (27.6 °C, ±1.2 SD); however, this difference was not statistically significant (*p* = 0.06). Minimum growth temperature (Tmin) and niche width (Tmax – Tmin) did not significantly differ between the two groups (*p* = 0.91). In addition to differences in growth observed using *in vivo* fluorescence measurements, we compared one warm and one cool temperature isolate (LA127 and LA31 respectively) when temperature was increased from 22 to 28 °C, closer to their Topt (Table S2). This showed that the warm-temperature isolate accumulated ~2× more volume-normalized particulate organic carbon (POC; *p* = 0.002; Figure S2A) and maintained a higher, but not statistically significant (*p* = 0.07), growth rate (Figure S2B).

**Table 1:** Traits derived from thermal performance curves (TPC) characterized for 11 *Synechococcus* isolates (indicated with n). TPC parameters are reported in °C.

| Temperature | n | Parameter | Mean | SD | Max | Min |
| --- | --- | --- | --- | --- | --- | --- |
| **22° (Initial)** | 1 | Tmax | 33.8 |  |  |  |
|  |  | Topt | 27.9 |  |  |  |
|  |  | Tmin | 11.2 |  |  |  |
|  |  | Width | 21.8 |  |  |  |
| **18°** | 5 | Tmax | 33.4 | 1.0 | 34.3 | 31.7 |
|  |  | Topt | 27.6 | 1.4 | 29.8 | 26 |
|  |  | Tmin | 12.4 | 3.1 | 14.8 | 9.0 |
|  |  | Width | 21.0 | 2.5 | 24.5 | 18.9 |
| **30°** | 5 | Tmax | 35.6 | 0.5 | 36.5 | 35.3 |
|  |  | Topt | 29.8 | 1.8 | 31.3 | 26.8 |
|  |  | Tmin | 14.2 | 4.4 | 19.1 | 9.0 |
|  |  | Width | 21.4 | 4.7 | 26.3 | 16.1 |

**Figure 1:**
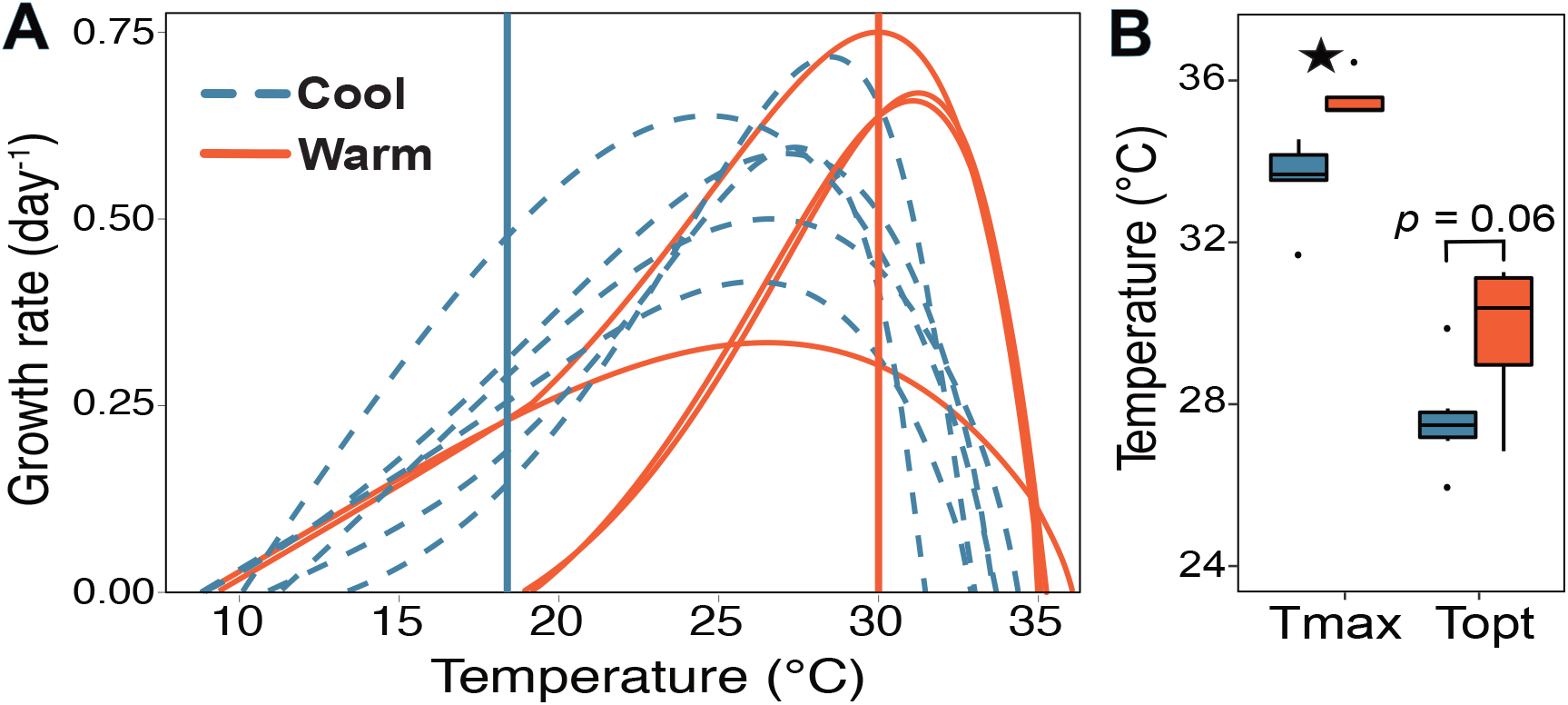
Thermal growth rate responses of *Synechococcus* isolates examined in this study, depending on whether the isolate came from cool (blue; 18-22 °C) or warm (red; 30 °C) incubation experimental temperatures. **A** Thermal Performance Curves (tpc) for all isolates determined using the Eppley-Norberg approach. Vertical lines indicate 18 °C (blue) and 30 °C (red). **B** Boxplots showing the maximum temperature limit (Tmax) and optimal temperature (Topt) for the two sets of isolates. Error bars represent quartiles, and the star indicates *p* < 0.05 level (t-test).

To compare genomic differences between cool and warm temperature isolates, sequence data for each isolate collected from this study (n=11) was assembled and manually curated producing estimated 100% complete draft genomes (Table S3). Recovered short read genomes (hereafter called draft assemblies) were 2.74 Mbp long (±0.01 SD), split between an average of 22.1 (±6 SD) contigs with a mean GC content of 63.3% (±0.00) and mean gene count of 2976.0 (±11 SD) for each isolate. We generated long reads for one cool (LA31) and one warm temperature isolate (LA127) and were able to close the genome of the warm temperature isolate (one contig 2.75 Mbp long compared to 27 contigs from the short-read only assembly). Including long reads substantially improved the assembly of the cool temperature isolate, reducing the number of contigs from 18 to six (Table S3).

When constructing a phylogenetic tree with all presently available *Synechococcus* genomes (Figure S2), the isolates from this study fell within the same clade as marine subcluster 5.2. With the level of resolution provided by 239 concatenated amino acid sequences, all 11 were nearly indistinguishable from each other (Figure 2A). Interestingly, the most closely related isolate in Genbank was CB0101 from Chesapeake Bay, another large coastal estuary located on the east coast of the United States (24). High genetic relatedness was even more apparent using average nucleotide identity (ANI), with greater than 99.99% (±0.003 SD) across 98.6-100% of the assembly for all genomes (Figure 2B). For comparison, the average ANI between isolates from this study and CB0101 was 85.46%.

**Figure 2:**
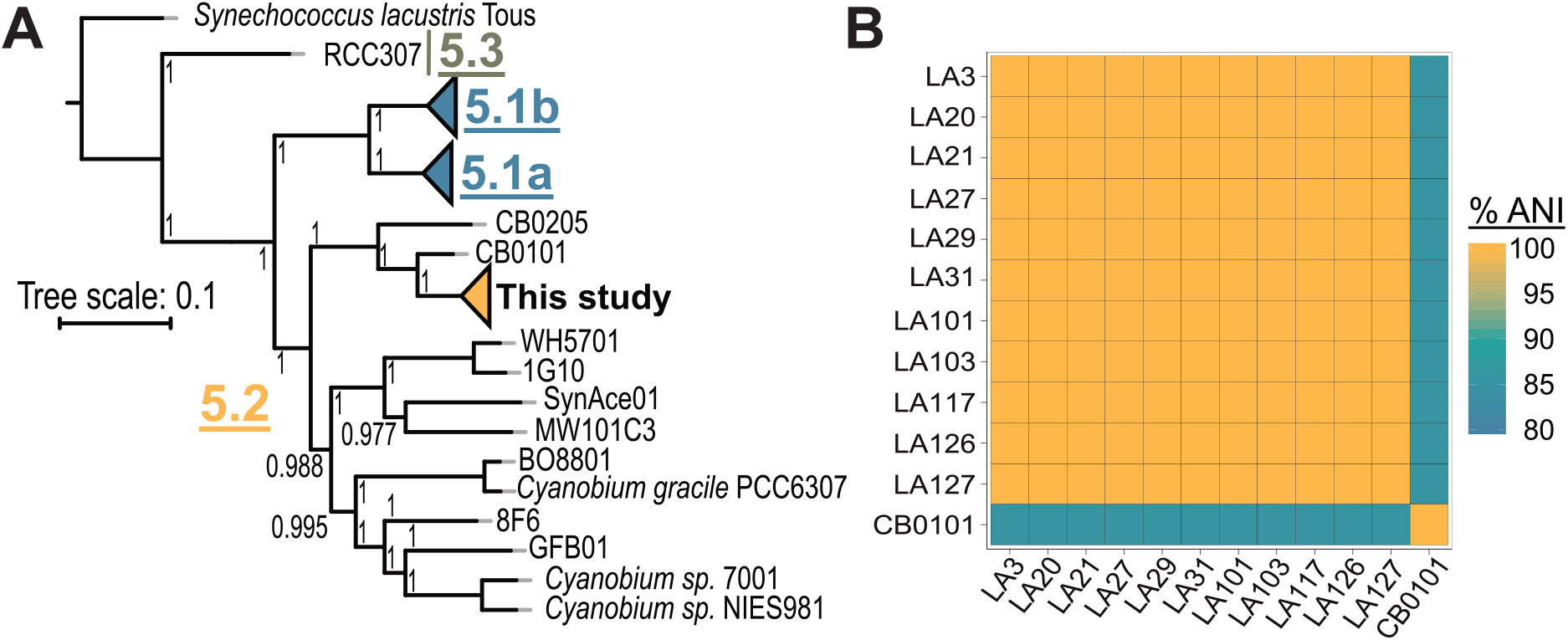
**A** Maximum likelihood tree showing relatedness of the three subclusters (5.1-5.3) comprising marine *Synechococcus*, using *Synechococcus lacustris* Tous to root the tree. Phylogenies were established by concatenating AA sequences of 239 single copy core genes. Scale bar shows AA change per position. **B** Average nucleotide identity (ANI) between all 11 Narragansett Bay isolates from this study. CB0101 from the Chesapeake Bay is the most closely-related genome present in Genbank and included for comparison.

Despite the high degree of similarity, when comparing draft assemblies from a pangenomic perspective, 62 out of 2985 gene clusters were identified by Anvi’o (Figure S4, Table S4) as having less than 100% functional (e.g. differences in sequence) or structural (e.g. insertions, deletions) homogeneity between one or more assemblies. Manually examining these gene clusters for genomic differences that correlated with isolation temperatures and measured thermal performance curves revealed that many of the features causing this detected variation were typically found only in one or a few isolates (Figure S5).

Only two gene clusters examined had patterns of variation that correlated with the temperature treatments that these strains were isolated from (e.g. exclusive to warm temperature isolates). These two gene clusters contain genes coding for the α and β subunits of the photosynthetic accessory pigment C-phycocyanin (*cpcA* and *cpcB* respectively). Cool temperature assemblies contained a complete copy of *cpcA*, while warm temperature assemblies lacked a complete copy of the α phycocyanin subunit (Figure S6A). On the other hand, all but one warm temperature isolate (LA126) had a complete copy of *cpcB*, while assemblies from cool temperature isolates only had the first 25 and the last 56 amino acids from *cpcB* coded for on different contigs (Figure S6B).

Because these differences in assembly between cool and warm temperature isolates are for genes coding for accessory pigments, we compared photosystem function between phenotypically distinct warm and cool temperature isolates. We measured whole-cell fluorescence spectra matching C-phycocyanin across rising temperatures for cool temperature strain LA31 (Topt = 27.5 °C, Tmax = 31.7 °C) and warm temperature strain LA127 (Topt = 29.7 °C, Tmax = 35.3 °C), using increasing fluorescence from initial levels at 22 °C as an indicator of loss of photosynthetic function to heat stress. The warm temperature isolate had a lower change in fluorescence at physiologically-relevant temperatures below 45 °C (Figure 3A). Fluorescence for both isolates increased exponentially between 48-54 °C, before abruptly crashing to zero at 57 °C. This suggests that light-harvesting energetic losses to fluorescence increase as the photosynthetic antenna complex becomes stressed by warming temperatures, before completely disassociating at a critical high temperature and losing all fluorescence. The warm temperature isolate had lower fluorescence at the fluorescence peak 54 °C (t-test, *p* = 0.07), suggesting it is better able to maintain its functionality under extreme thermal stress (Figure 3A). Furthermore, both isolates had differing photophysiologies when comparing the photosynthetic efficiency parameter Fv/Fm (Figure 3B). When acclimated to 28° C the warm temperature isolate had significantly higher values (t-test, *p* = 1.04 × 10^−7^), suggesting its photosystem II had a greater photochemical efficiency than the cool temperature isolate. The values reported here are analogous to Fv/Fm measurements reported for other marine *Synechococcus* spp. (25).

**Figure 3:**
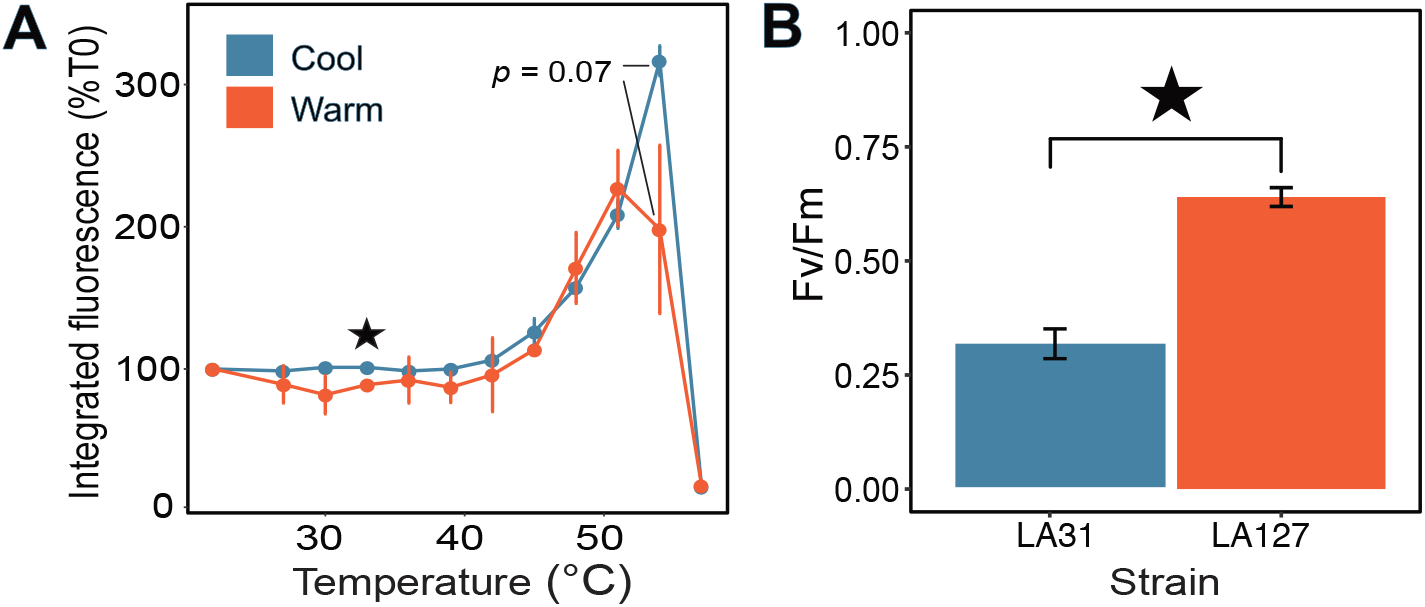
**A** Mean whole-cell integrated fluorescence (600-700nm) with temperature for warm and cool strains and (**B)** Photochemical efficiency (Fv/Fm) of PSII at an excitation wavelength matching phycocyanin and allophycocyanin (645 nm). Blue colors indicate the cool temperature (18-22 °C) isolate LA31, while red shows warm temperature (30 °C) isolate LA127. The star indicates observations that were statistically different (t-test, *p* < 0.05), and error bars represent ± 1 SD from triplicate trials.

A closer look at the genes associated with C-phycocyanin was unable to discern the exact genetic mechanism causing these measured differences in photosynthetic function. We compared the closed warm temperature isolate genome (LA127) and the closed genome of the closely related Chesapeake Bay strain CB0101, which revealed the majority of C-phycocyanin and surrounding genes in the same orientation (Figure 4A). In all draft assemblies (short-read data only), assembly failed at this exact locus (Figure 4B & C). Interestingly however, the pattern of contig breakage is conserved for all draft assemblies of isolates from cool temperatures (breaking within copies of *cpcB*, Figure 4B), while a distinct pattern of contig breaks is found in assemblies of warm temperature isolates (breaking sooner in all cases, Figure 4C). These conserved contig-breakage patterns within our cool and warm groups correlate with their measured differences in thermal performance curves and photophysiology, suggesting whatever variation is consistently leading to these assembly results may be involved in these phenotypic traits. Long-read sequencing and assembly of all isolates may help further interrogate this region. For the two we were able to attempt, although we spanned this difficult-to-assemble region for the warm temperature isolate LA127, we were not able to for the cool temperature isolate LA31 (Figure 4D), preventing a direct comparison.

**Figure 4:**
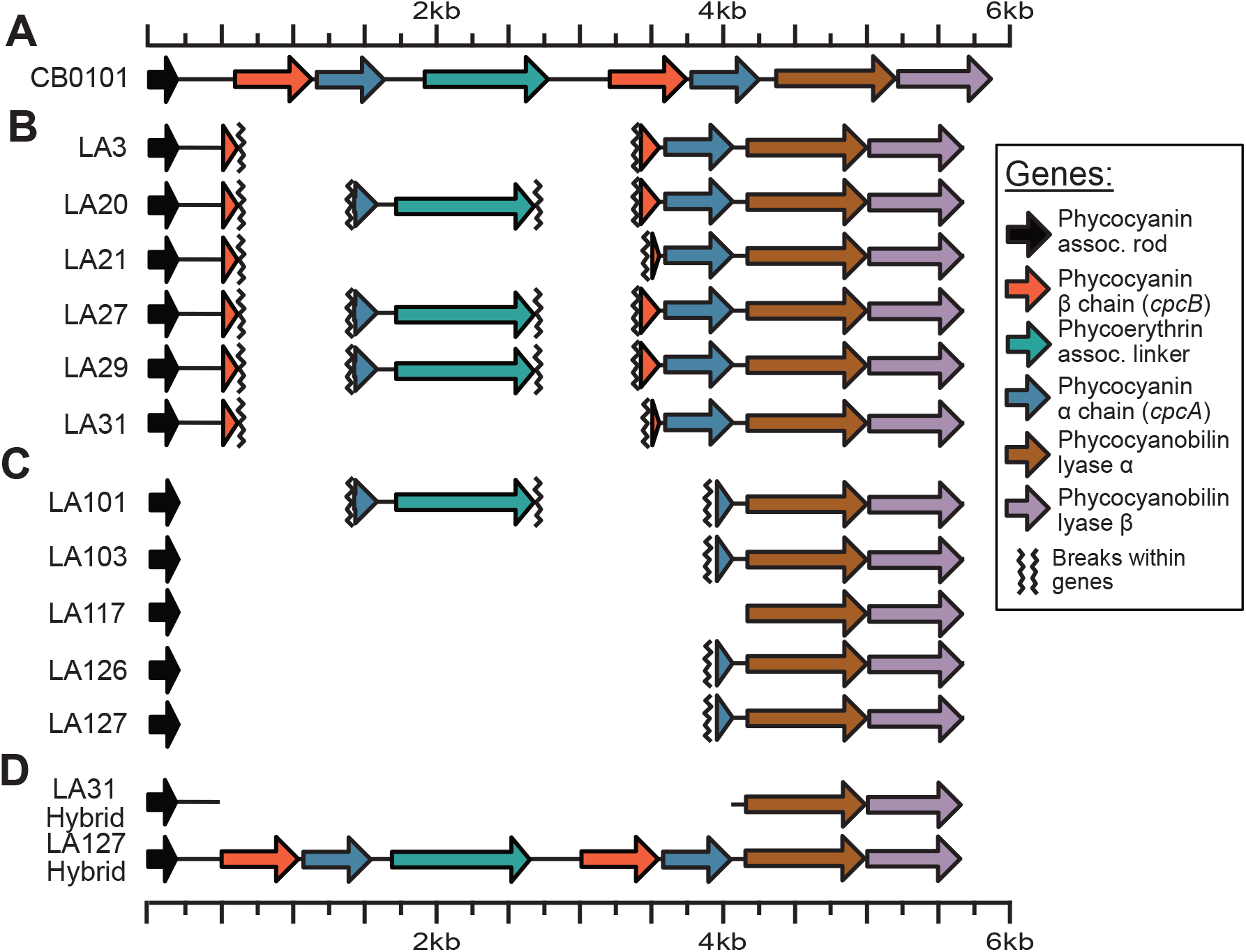
Assembly and coverage information of genes within the locus containing the primary C-phycocyanin genes, *cpcA* (blue arrows) and *cpcB* (orange arrows). **A** C-Phycocyanin locus in the closest related genome available on NCBI, CB0101. **B** Structure of draft assemblies of isolates recovered from 18° or 22° **(C)** and 30° C (**D).** Same locus in a cool and warm temperature strain incorporating long reads to close assembly gaps in a hybrid assembly approach.

Mapping Illumina short-reads from all isolates to single complete copies of *cpcA* and *cpcB* from draft assemblies showed some single nucleotide variants (SNVs; Supplemental Note 1 and Figure S7); however, no SNVs were detected when mapping short-reads to this locus on the closed genome from LA127 (Supplemental Note 1 and Figure S8). This suggests that although minor sequence differences were detected between the complete copies of these genes recovered in the closed hybrid genome (Table S5 & S6), they were identical to the completely assembled copies in the draft assemblies (Table S7). Mapping rates were used to attempt to detect different gene copy numbers at this locus which might contribute to this systematic pattern contig breakage (Supplemental Note 2); however, these data were inconclusive. Copy numbers of *cpcA* and *cpcB* vary across Subcluster 5.2 genomes (Table S8) and these isolates appear to have multiple copies of the genes (Figures S7-S10; Supplemental Note 2); however, we were not able to deduce from these data whether or not copy number differed between isolates from different temperatures. Although the exact genomic differences causing these assembly results are as yet unclear, these contig breaks occur systematically between cool and warm temperature isolates. As this correlates with the observed temperature responses (Figure 1), it suggests that genes associated with accessory pigment C-phycocyanin production could be involved in thermal adaptation.

## Discussion

This study demonstrates the coexistence of distinct thermal phenotypes within a highly genetically similar, single population of coastal cyanobacteria. The division of this estuarine *Synechococcus* population into contrasting cool and warm thermotypes was revealed following incubation experiments at 18 °C and 30 °C, in which strain-sorting was identified by culture isolations and thermal phenotype determinations in the laboratory. By using high-coverage short reads as well as long reads to reconstruct these isolates' genomes, we found that this striking phenotypic divergence between thermotypes appears to be tied to very minor genetic differences. Although previous work has established that distinct functionally-relevant ecotypes coexist within populations of picocyanobacteria (e.g. Kashtan et al., 2014; Thompson & Kouba, 2019), this is the first study to report coexisting ecotypes at this high level of genomic resolution.

Variation in assembly of the genomic region containing genes coding for C-phycocyanin correlated with isolation temperature and measured thermal phenotype. In addition, we measured distinct differences in both thermal resilience of the phycobilisome accessory pigment complex and photosynthetic efficiency (Fv/Fm) between the two thermal ecotypes isolated from different temperatures. These genomic and photophysiological differences are consistent with previous work examining thermal adaptation in marine *Synechococcus*. For instance, it has been suggested that differences in light-harvesting machinery can explain the global distribution of *Synechococcus* clades across large temperature differences (27, 28), and variation in a single amino acid in either the α and β units of R-phycocyanin, an ortholog of C-phycocyanin, correlated with *Synechococcus* thermal adaptation (29). Pittera et al. (2017) also found that at elevated temperatures a tropical, low-latitude strain had lower fluorescence of the antenna pigment complex (indicating more efficient photosynthetic energy capture) than a sub-polar strain. This is similar to the trend we observed between warm and cool temperature phenotypes in our study (29). Further supporting the role of phycocyanin in *Synechococcus* thermal adaptation, it has been observed that intracellular concentrations of phycobilisome proteins increase under high temperatures (25). It should also be noted that there could be additional, non-genomic factors such as epigenetic effects not tested for in this study that can increase phenotypic heterogeneity within a population, even at a high degree of relatedness. For instance, epigenetic differences have sometimes been found to be associated with bacterial stress responses (30, 31).

Although our experimental setup precludes looking at the relative abundance of each ecotype in the original population, we note that the one isolate collected directly from the environment was the cool temperature ecotype. In an environmental context, the average summertime surface water temperature at our sample site for the period from 1957 to 2019 was 20.6 °C, with a maximum of 26.5 °C (Figure 5A). This distribution of temperatures falls below the Topt for both ecotypes, but would likely favor cool temperature isolates. Warm temperature isolates were also collected from 30 °C enrichments, 3.5 °C above the highest measured temperature at this site. Average summer SST at this site has been increasing at a rate of 0.03 °C per year since 1957 (Figure 5A), meaning that the average summertime SST in Narragansett Bay will likely increase to ~23 °C by the end of the century. Assuming a similar distribution of temperatures in the year 2100, there will be periods when SST is above the average Topt of the low temperature ecotype, and conditions will favor the high temperature ecotype (Figure 5B). Any continued trend of rising temperatures beyond 2100 will continue to further expand the niche of the warm temperature ecotype.

**Figure 5:**
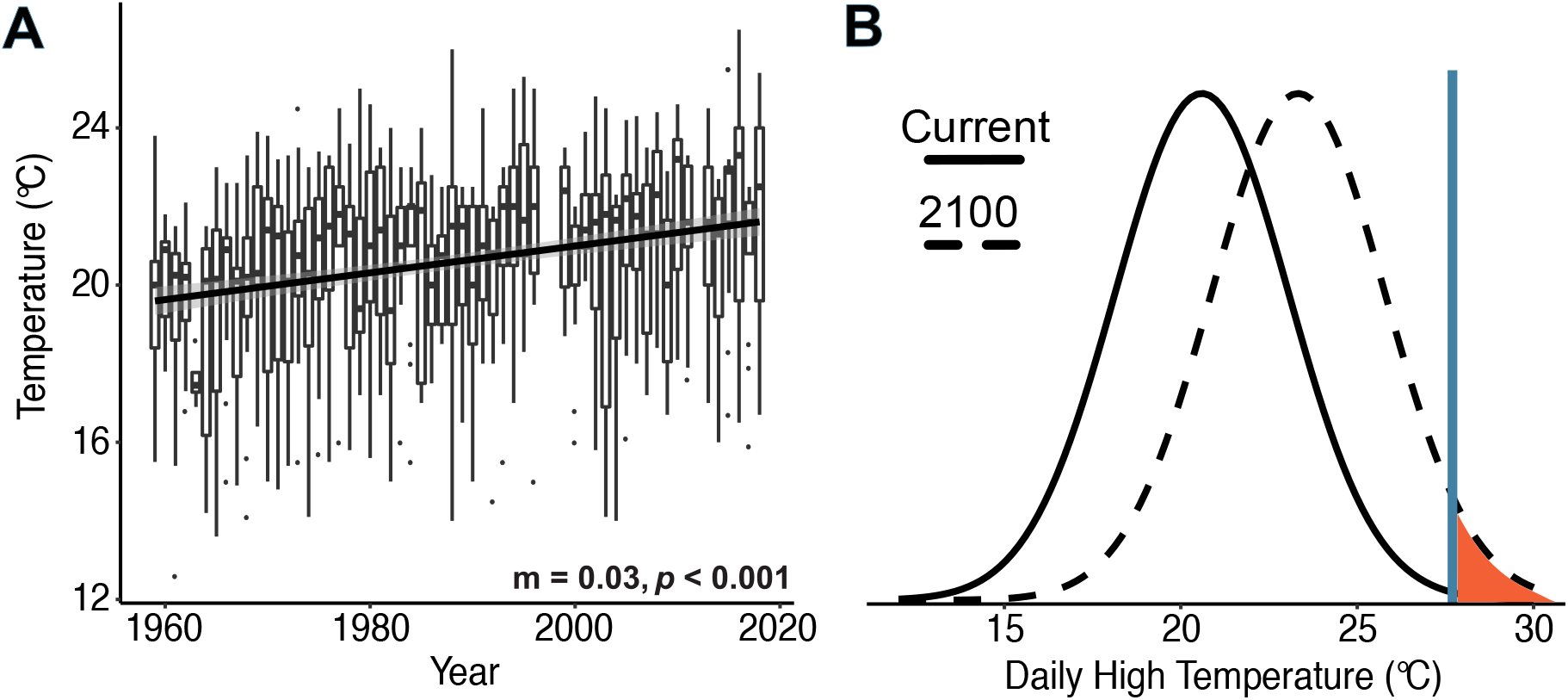
**A** Boxplot of summertime sea surface temperature (SST) increases at the Narragansett Bay Time Series from 1957 to 2019. Trendline shows the output of a linear model fit to the data. The slope of this model and the *p* value are shown below the data. **B** Hypothetical normal seasonal temperature distributions created using the mean of the recent data shown in panel **A** (solid line), and the predicted distribution of these data in the year 2100 (dashed line) using the slope of the linear model. A blue vertical line shows the average Topt for all cool temperature isolates. Temperatures above this line, which will likely favor warm temperature ecotypes, are shown in red.

In addition to this intraspecific diversity likely increasing the resilience of this population to long term warming trends, it is also interesting to consider why adaptations to temperatures exceeding current thermal maxima are maintained in this population. The higher growth rates of cool temperature isolates under typical summer conditions suggest that having the warm temperature phenotype has a fitness cost (when defined purely by growth rates), and in theory selection (considering only growth rates) should remove this phenotype from the population (Innan & Kondrashov, 2010 and references therein). Given the temperature trends at this site, it seems unlikely these are seasonal ecotypes, as the shape of thermal curves suggest that the warm temperature phenotype only has a growth advantage above the maximum daily high temperature observed (26.5 °C).

An intriguing explanation for this thermal diversity is that these microbes originated in warmer low latitude waters, and were advected into this relatively cooler region as part of the northerly flow of the nearby Gulf Stream. It has been estimated that microbes entrained in the Gulf Stream may experience a range of temperatures as a result of advection that is larger than changes due to seasonal patterns (33). Although Narragansett Bay is a narrow coastal estuary, wind-driven circulation during summer months facilitates persistent exchange between estuarine waters in the Bay and oceanic waters in Rhode Island Sound (34). This has led to the conjecture that allochthonous inputs of sub-tropical phytoplankton could occur (35, 36), which is supported by the well documented recurring appearance of subtropical fish species in Narragansett Bay during the summer (37).

These coexisting temperature phenotypes are also interesting in the context of marine *Synechococcus* evolution, as temperature is thought to be a key driver of diversity between clades within this group (8, 27). It is possible that intraspecific microdiversity of thermal phenotypes could have been a contributing mechanism in the diversification of *Synechococcus* into the distinct lineages observed today. When a population consisting of multiple thermotypes encounters a novel thermal environment, one phenotype may be selected over another, potentially leading to genetic divergence and speciation. Further studies will be needed on intraspecific microdiversity at this level, between nearly identical strains, to assess its potential role in the evolution of *Synechococcus* and marine microbes in general.

Our findings also have implications for our general understanding of biological responses to rising temperatures. It has been shown that there can be a greater diversity of responses to ocean acidification between ecotypes within phytoplankton taxonomic and functional groups, than between them (Schaum et al. 2013, Hutchins et al. 2013). Our findings show that ecotypes with distinct responses to climate warming can co-occur within a single population. This microdiversity in thermal traits also has been detected in other marine phytoplankton. In a similar study conducted within Narragansett Bay, thermal performance curves of recently isolated strains of the diatom genus *Skeletonema* were compared and showed a similar high degree of intraspecific diversity of thermal traits (39). This study also observed a similar significant difference in thermal maxima (Tmax) across strains, and suggested that variability at such thermal limits plays an important role in both ecological and biogeochemical dynamics. Because of the high degree of genetic similarity between these isolates, amplicon sequencing or metagenomic microbial surveys would not be able to detect this level of functional microdiversity.

Taking together, the findings of the current study and those of the aforementioned diatom study (39) suggests that this type of fine-scale variation may be widespread among marine microbial taxa. In the case of our estuarine *Synechococcus*, this cryptic thermal microdiversity will likely contribute to this population’s ability to continue occupying its picoplanktonic niche even in the face of considerable increases in environmental temperatures. Another important implication is that culture studies using a single isolate or strain from a population may underestimate that population’s resilience to warming. A better understanding of the existing functional thermal diversity within populations is needed to correctly model the impact that future elevated temperatures will have on microbial communities, and on the biogeochemical cycles that they regulate.

## Methods

### Sampling and Cell Isolation

Surface water at a temperature of 22°C and salinity of 28.48 was collected from the Narragansett Bay Time Series site (latitude 41.47, longitude −71.40) on July 18^th^, 2017 (36). Collected surface water was pre-filtered using 200μm mesh to remove debris and large grazers. In order to select for multiple temperature phenotypes, we split the collected seawater into 18°, 22° (control), 26°, and 30° C temperature treatments. Incubations were performed in triplicate 2L polycarbonate bottles under a 12:12 light dark cycle at 150 μmoles photons / m^2^ * sec^−1^. Cultures were amended with nutrients to match F/40 media (40), and diluted semi-continuously with 0.2 um-filtered seawater medium when chlorophyll a fluorescence reached a predetermined threshold to prevent nutrient depletion and avoid cells entering stationary phase.

At the time of seawater collection and after 10 days, cells from all temperature treatments that were <1.5 μm in diameter with measurable phycocyanin fluorescence were sorted into 96-well plates containing F/20 media (40) using a BD Influx (San Jose, CA, USA). Wells showing growth over time were transferred into artificial seawater (Sunda, Price, & Morel, 2005), and nutrient concentrations were gradually adjusted to F/2 levels (40). All isolates were maintained at 22° C at a light intensity of 150 μmoles photons / m^2^ * sec^−1^, with weekly transfers into fresh culture medium.

### Thermal Performance Assays

Thermal performance curves were obtained from 11 strains sorted from the initial surface seawater and from the 18 and 30 °C treatments (Table 1). This was done by acclimating aliquots of each culture for two weeks to temperatures between 9° and 33°. Temperatures >33° were added where permissible. This temperature range was chosen because it exceeds that of Narragansett Bay (0.5-24.6 °C, Rynearson, Flickinger, & Fontaine, 2020), and encompasses projected SST increases (20). Strains were grown at each temperature in triplicate 8 ml borosilicate vials containing 5ml of F/2 medium with a 12:12 light:dark cycle and 150 μmoles photons / m^2^ * sec^−1^. Biomass was recorded every two days using *in vivo* chlorophyll a fluorescence measured on a Turner AU-10 fluorometer (Turner Designs Inc., Sunnyvale, CA, USA), and growth rates and Eppley-Norberg thermal performance curves (43) were calculated in R (R Team, 2019) using the package growthTools (DOI:10.5281/zenodo.3634918). Cultures containing algal contaminants (verified using fluorescence microscopy) were excluded from the dataset. In two strains, LA20 and LA27, after two weeks of acclimation at 9 °C no *Synechococcus* cells were observed in the culture, so the growth rate was set to zero for these cultures. The thermal performance curve of each strain is hereafter referred to as its phenotype.

We verified growth rates for two strains, LA31 and LA127, isolated from low (18 °C) and high (30 °C) temperature treatments respectively using changes in particulate organic carbon (POC). Strains were grown in triplicate in 1L polycarbonate bottles for two weeks at 22 °C (12:12 light:dark cycle and 150 μmoles photons / m^2^ * sec^−1^) with dilutions every three days. In addition to POC, carbon fixation was also measured for both strains. Analysis for both POC and carbon fixation were done as in Qu, Fu, & Hutchins (2018) and references therein. At the end of two weeks, cultures were diluted to equal biomass and the temperature was increased to 28 °C. Both isolates were sampled at the beginning, after two days, and after four days.

### DNA Extraction, Sequencing, and Analysis

250ml of stationary phase culture for each strain was filtered onto 0.2 μm Polyethersulfone (PES) membrane filters and DNA was extracted using the DNeasy PowerSoil kit (Qiagen, Germantown, MD, USA). Sequencing was done on an Illumina Hiseq, paired-end with 150 basepair reads (2×150) with ~10 million reads per sample at Novogene Inc. (Beijing, China). The quality of base calls was assessed using fastqc (https://www.bioinformatics.babraham.ac.uk/projects/fastqc), and reads were assembled with SPAdes version 3.13 (46). All programs were run with default settings, unless otherwise noted. To assist in genome curation, reads were assigned taxonomy using Centrifuge version 1.0.4 (47) trained on the “p_compressed+h+v” index (downloaded July 2019), and raw reads were mapped to their assembled genomes using bowtie2 version 2.3.5 (48). The resulting assemblies were curated to remove associated heterotrophs using Anvi’o version 5.5 (49) based on tetranucleotide frequency, coverage, and read taxonomy. Gene calls for curated assemblies were generated using Prodigal version 2.6.3 (50) which were then annotated using kofamScan version 1.1.0 (51) and imported into Anvi’o.

In addition to the short-read Illumina sequencing, DNA was extracted from low-temperature (18 °C) strain LA31 and high-temperature (30 °C) strain LA127 for long-read sequencing using an Oxford Nanopore Minion (Oxford, UK) with the FLO-MIN106D flow cell. Library prep was done using the Ligation Sequencing Kit (SQK-LSK109) and Rapid Barcoding Kit (SQK-RBK004) following the Genomic DNA by Ligation protocol (https://store.nanoporetech.com/us/media/wysiwyg/pdfs/SQK-LSK109/Genomic_DNA_by_Ligation_SQK-LSK109_-minion.pdf). 200ml of culture grown to stationary phase was concentrated using centrifugation (27,000 × g for 15 minutes) and extracted with the GenElute Bacterial Genomic DNA Kit (Millipore Sigma, Burlington, MA, USA). Basecalling was done using Guppy version 2.2.3 and long reads were filtered using filtLong version 0.2.0 (https://github.com/rrwick/Filtlong). Filtered long reads were mapped to their respective draft assemblies using Minimap2 version 2.17 (52), and the same was done for the short reads using bowtie2 (48). Mapped long and short reads were then assembled together using Unicycler version 0.4.8 (53) with subsequent gene calling and annotation performed as described above.

In order to place our isolates in the context of the broader diversity of *Synechococcus*, we pulled all *Synechococcus* genomes (a total of 78 as of July 2019) from NCBI’s Refseq (Pruitt & Maglott, 2001; Table S9) and placed them on a phylogenetic tree constructed with GToTree version 1.4.11 (55) using concatenated amino acid sequences for 239 single copy core genes specific to cyanobacteria (using the “Cyanobacteria.hmm” included within GToTree). In short, genes were identified with HMMER3 version 3.2.1 (56), aligned with muscle version 3.8.1551 (57), trimmed with trimal 1.4 (58), and concatenated before calculating phylogenetic distance using FastTree2 version 2.1.10 (59). All trees were visualized using the interactive Tree of Life webpage (60). Average nucleotide identity (ANI) was calculated using fastANI version 1.2 (61). The Anvi’o pangenomic pipeline (62) was used to identify gene clusters and to test for genecluster correlations with the original incubation temperature from which each strain was isolated. In addition, we looked for sequence variants at loci of interest by mapping reads to the nearest phylogenetic neighbor with a complete genome and profiling single-codon-variants (SCV), utilizing the framework available within Anvi’o (demonstrated at http://merenlab.org/2015/07/20/analyzing-variability/).

### Photophysiology Measurements

To analyze differences in photosynthetic accessory pigment function, 200ml of triplicate cultures of low-temperature (18 °C) strain LA31 and high-temperature (30 °C) strain LA127 grown to stationary phase at 22 °C were concentrated by centrifuging for 15 minutes at 27,000 × g. Cell pellets were then resuspended in 5ml of sterile media, and the fluorescence and absorption spectra measured on a SpectraMax m2^e^ (Molecular Devices, San Jose, CA, USA). In order to detect changes in efficiency in the light gathering mechanisms with temperature, fluorescence was measured every three degrees from 22°-57° (10 minute incubation at each temperature) following the methods of Pittera, Partensky, & Six, (2017). This large range of temperatures was used in order to detect the instantaneous disassociation temperature of the phycocyanin antenna complex. Fluorescence emission was measured from 600-700nm (530nm excitation wavelength) matching the profile of the allophycocyanin/phycocyanin pigment complex (63). In addition, we measured the photosynthetic efficiency of photosystem II (Fv/Fm) for these two strains when acclimated to 28° C using a PHYTO-PAM with an excitation wavelength set to 645 nm for C-phycocyanin and allophycocyanin (Heinz Walz, Effeltrich, Germany). Fv/Fm measurements were made using triplicate cultures and three technical replicates each that were dark acclimated for 20 minutes, as in McParland et al. (2019).

### Analysis of Long-term Temperature Trends

In order to explore changes in summertime temperature trends at our study site, all available SST data from 1957 through 2019 collected as part of the long-term times series weekly measurements was downloaded from https://web.uri.edu/gso/research/plankton/data/. All measurements from June-August were aggregated by year and a simple linear model (using the lm command in R) used to calculate the rate temperature increase. The slope of this linear model was then used to predict the distribution of summertime temperature in the year 2100. A normal distribution was assumed for both present day and future temperatures, as well as a similar standard deviation from the mean.

## Data and Code Availability

Curated genomes are available from SRA under the BioProject ID PRJNA566206. Isolate information and individual BioSample accession numbers can be found in Supplemental Table 2. Scripts used in the analysis and generation of all figures as well as all physiological data are available at https://figshare.com/projects/Kling_et_al_2020/66188. Phenotypic data are also available at: www.bco-dmo.org/award/712792.

## Acknowledgements

This work was funded by National Science Foundation Dimensions of Biodiversity grants OCE1638804 to DAH, OCE1638834 to TR, and OCE1638958 to EL, and by OCE 1851222 to DAH and EAW. This study is based upon work conducted at the URI Marine Science Research Facility supported in part by the National Science Foundation EPSCoR awards OIA-1004057 and OIA-1655221. Illumina sequencing was supported by the Global Climate Change and Ocean Atmosphere Interaction Project, Marine Biological Sample Museum Upgrade and Expansion Project (GASI-01-02-04) to QZ and CW. Thanks to David Kehoe (Indiana University) for helpful insights into *Synechococcus* photophysiology, and to J. Cameron Thrash (University of Southern California) and Ben Temperton (University of Exeter) for helping generate and perform base calling on Minion long reads.

## Notes

https://figshare.com/projects/Kling_et_al_2020/66188

### Competing Interest Statement

The authors have declared no competing interest.

