## Supplemental Tables for "Dual thermal ecotypes co-exist within a nearly genetically-identical population of the unicellular marine cyanobacterium *Synechococcus*"

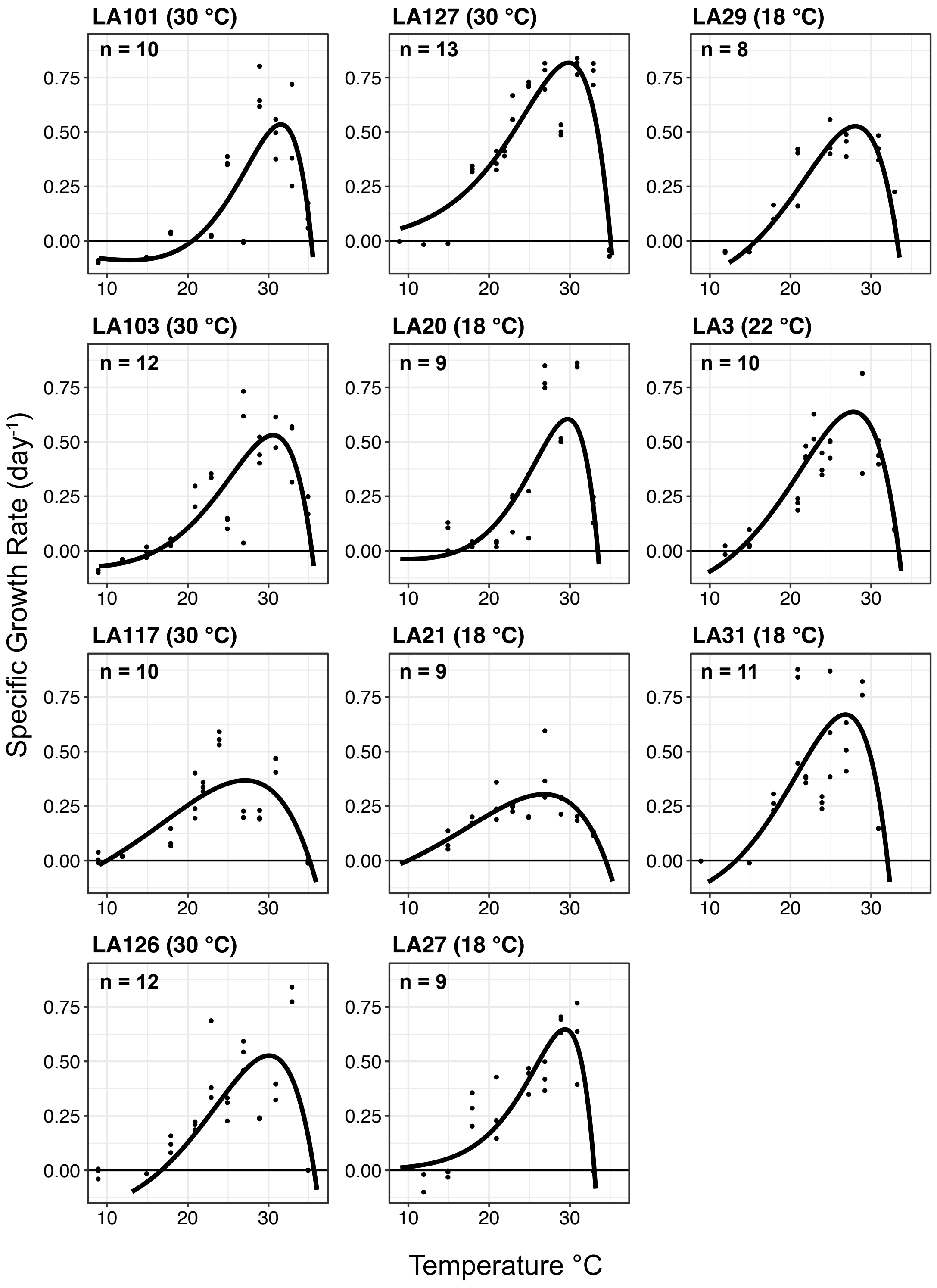


Figure S1: Thermal performance curves for 11 new *Synechococcus* isolates from Narragansett Bay, RI. Points indicate measured growth rates across temperatures while trend lines are the result of a Eppley-Norberg curve fit to the data.

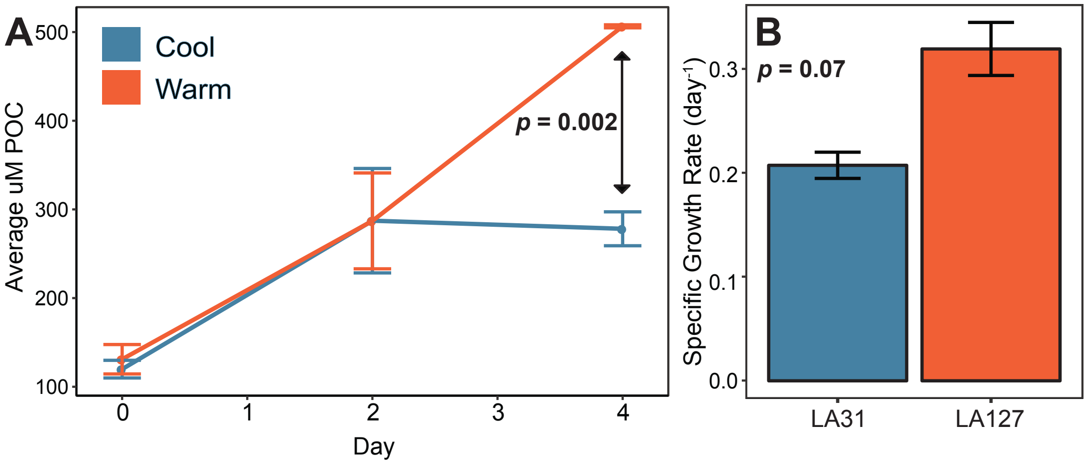


Figure S2: Large volume experiments with two isolate strains representing both cool (blue) and warm (orange) temperature isolates. **A** Accumulation of particulate organic carbon over the course of 4 days after increasing the temperature from 22 to 28 °C. **B** Growth rate from day 0 to day 4 for each strain derived from the change in POC. Error bars represent one standard deviation from the mean.

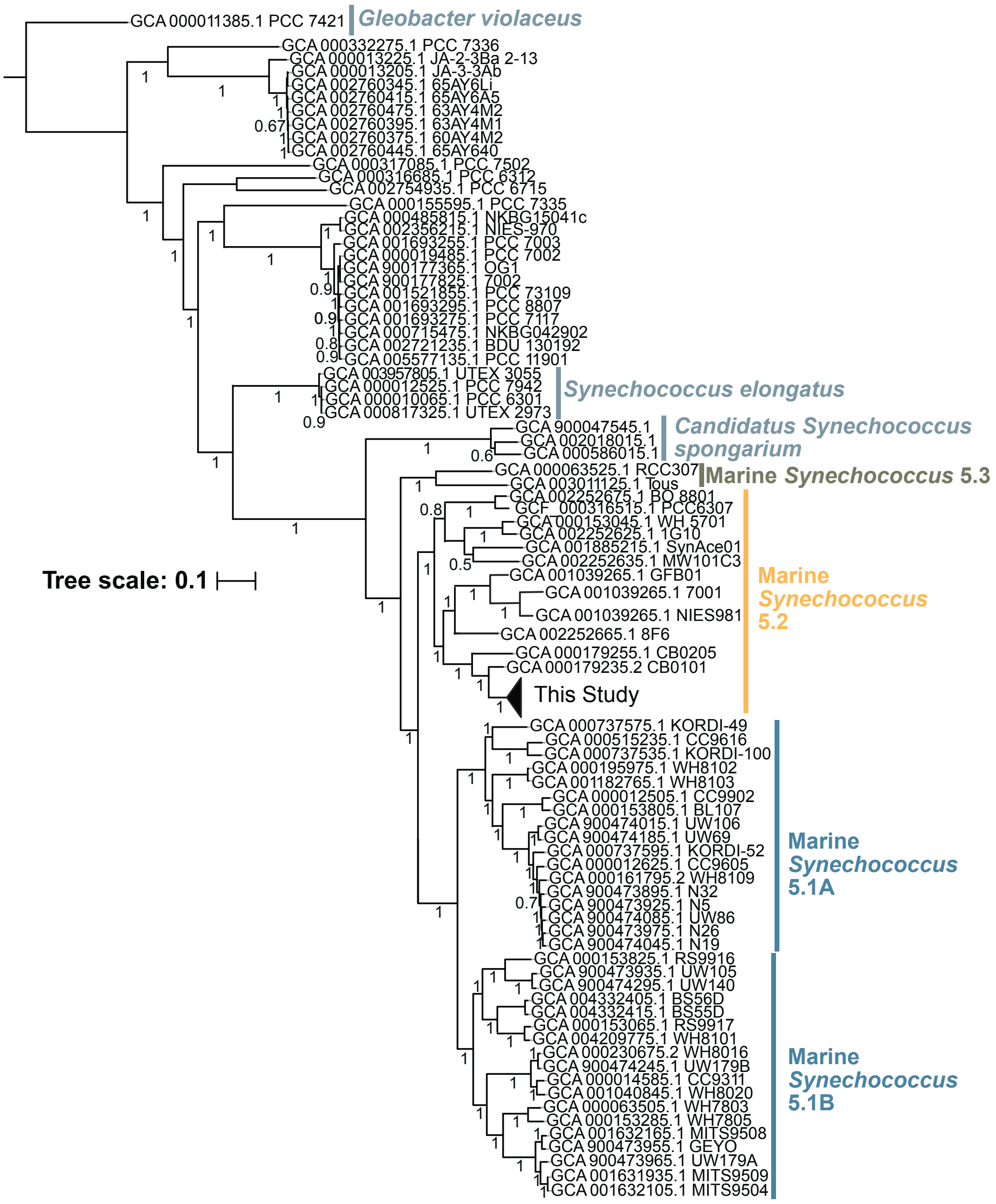


Figure S3: Approximate maximum-likelihood tree for all 11 isolates collected in this study and all 78 available complete *Synechococcus* assemblies in Genbank. *Gleobacter violaceus* is included as the outgroup. Based on concatenated amino acid alignments of 239 single-copy genes specific for Cyanobacteria. Scale indicates predicted amino acid changes per position.

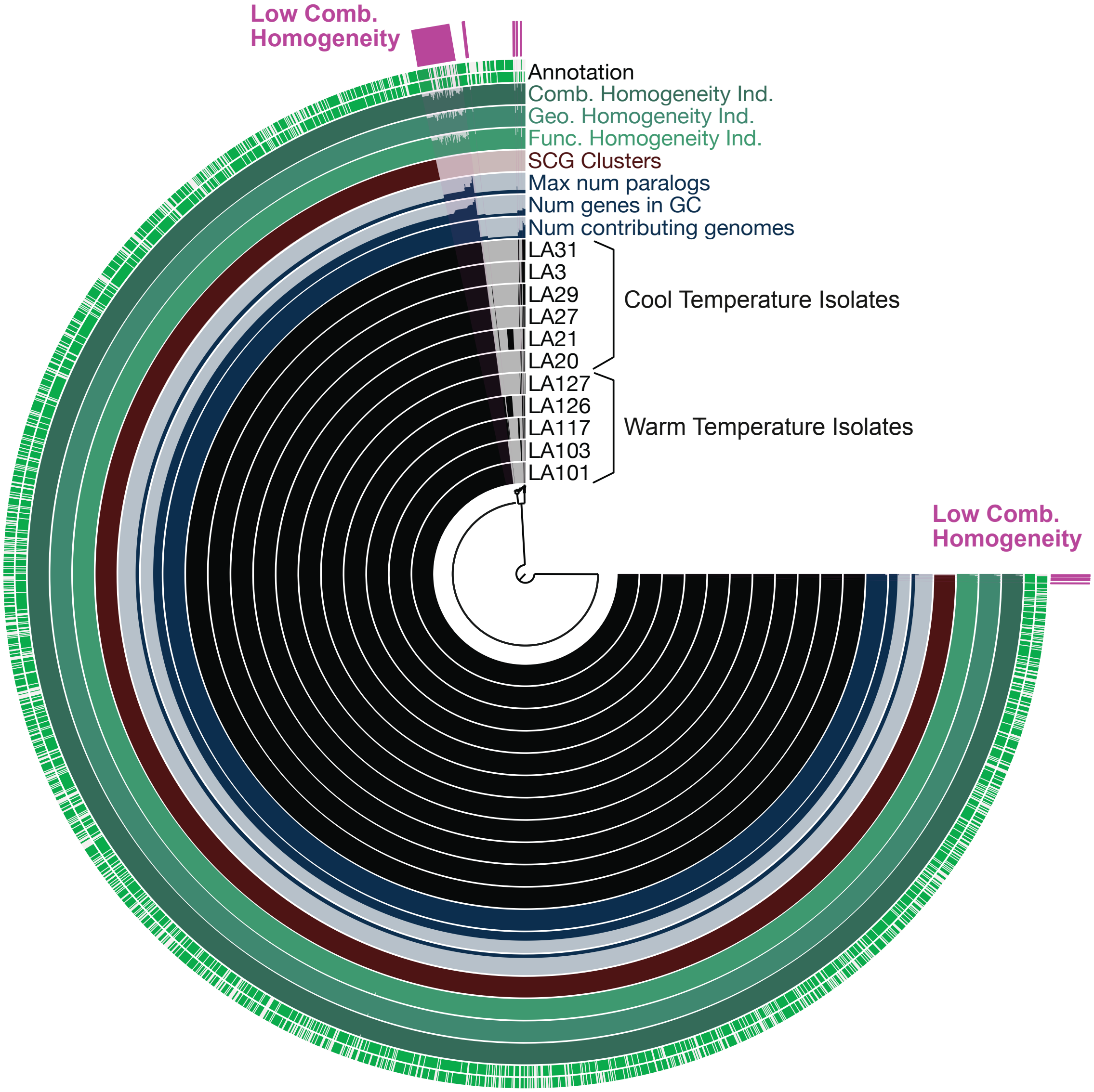


Figure S4: Visualization of Anvi'o pangenomic analysis. From inside to outside: tree represents tetranucleotide frequency in all 11 isolates analyzed in this study, with black regions showing the presence and absence of all gene clusters, number of genomes with a given gene cluster, number of genes in a given gene cluster, number of paralogs, number of single copy genes, homogeneity within a gene cluster, and successful annotation of the gene cluster (shown in green). Gene clusters with less than 100% homogeneity that were manually examined to detect genomic differences that correlate with temperature phenotype are highlighted with purple.

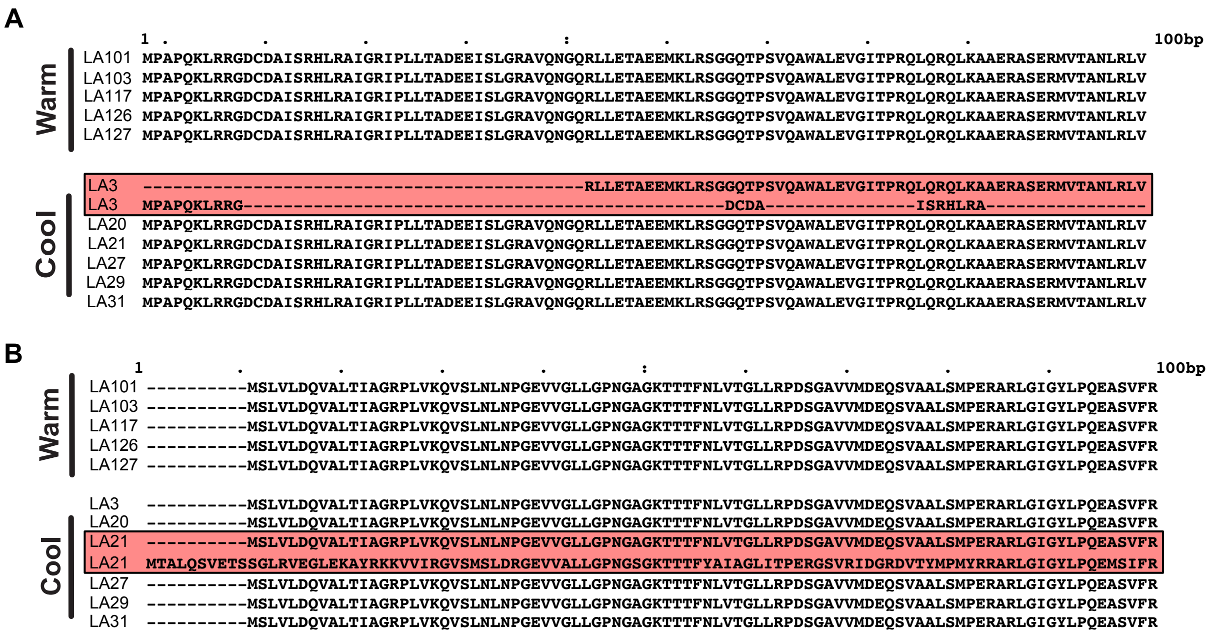

Figure S5: First 100 amino acids from two gene clusters identified by Anvi'o as having less than 100% homogeneity, but that were determined to not correlate with isolation temperature or the observed high/low temperature phenotypes: **A** GC0000045 shows the result of a poor alignment where one contig of the isolate LA3 has the 21 residues of this protein coded for on one contig that were mistakenly aligned along with a longer portion of that gene found on another contig. Likely, this is the result of a contig break within this coding region. **B** GC0000041 shows that a single isolate (LA21) has a duplicated version of this protein that is not shared by the other isolates.

**A**

```
                             1bp       .         .         .         .         :         .         .         .         .      100bp
Warm  LA101_cpcA_fragment    ----------------------------------------------------------------------------------------------------
      LA103_cpcA_fragment    ----------------------------------------------------------------------------------------------------
      LA126_cpcA_fragment    ----------------------------------------------------------------------------------------------------
      LA127_cpcA_fragment    ----------------------------------------------------------------------------------------------------

Cool  LA20_cpcA_complete     MKTPLTEAVAAADSQGRFLSNTELNAAFGRFERAANALAAAKALTAKADELVNGAAQAVYNKFPYTTQMQGANYASDARGKAKCARDIGYYLRQVTYCLV
      LA21_cpcA_complete     MKTPLTEAVAAADSQGRFLSNTELNAAFGRFERAANALAAAKALTAKADELVNGAAQAVYNKFPYTTQMQGANYASDARGKAKCARDIGYYLRQVTYCLV
      LA27_cpcA_complete     MKTPLTEAVAAADSQGRFLSNTELNAAFGRFERAANALAAAKALTAKADELVNGAAQAVYNKFPYTTQMQGANYASDARGKAKCARDIGYYLRQVTYCLV
      LA29_cpcA_complete     MKTPLTEAVAAADSQGRFLSNTELNAAFGRFERAANALAAAKALTAKADELVNGAAQAVYNKFPYTTQMQGANYASDARGKAKCARDIGYYLRQVTYCLV
      LA3_cpcA_complete      MKTPLTEAVAAADSQGRFLSNTELNAAFGRFERAANALAAAKALTAKADELVNGAAQAVYNKFPYTTQMQGANYASDARGKAKCARDIGYYLRQVTYCLV
      LA31_cpcA_complete     MKTPLTEAVAAADSQGRFLSNTELNAAFGRFERAANALAAAKALTAKADELVNGAAQAVYNKFPYTTQMQGANYASDARGKAKCARDIGYYLRQVTYCLV

                                       .         .         .         .         :       .162bp
Warm  LA101_cpcA_fragment    ------------------------SPSWYVEALNYIKANHGISGDAGVIANNYIDYAIAALV
      LA103_cpcA_fragment    ------------------------SPSWYVEALNYIKANHGISGDAGVIANNYIDYAIAALV
      LA126_cpcA_fragment    ------------------------SPSWYVEALNYIKANHGISGDAGVIANNYIDYAIAALV
      LA127_cpcA_fragment    ------------------------SPSWYVEALNYIKANHGISGDAGVIANNYIDYAIAALV

Cool  LA20_cpcA_complete     AGGTGPIDEYLIAGLDEINRAFELSPSWYVEALNYIKANHGISGDAGVIANNYIDYAIAALV
      LA21_cpcA_complete     AGGTGPIDEYLIAGLDEINRAFELSPSWYVEALNYIKANHGISGDAGVIANNYIDYAIAALV
      LA27_cpcA_complete     AGGTGPIDEYLIAGLDEINRAFELSPSWYVEALNYIKANHGISGDAGVIANNYIDYAIAALV
      LA29_cpcA_complete     AGGTGPIDEYLIAGLDEINRAFELSPSWYVEALNYIKANHGISGDAGVIANNYIDYAIAALV
      LA3_cpcA_complete      AGGTGPIDEYLIAGLDEINRAFELSPSWYVEALNYIKANHGISGDAGVIANNYIDYAIAALV
      LA31_cpcA_complete     AGGTGPIDEYLIAGLDEINRAFELSPSWYVEALNYIKANHGISGDAGVIANNYIDYAIAALV
```

**B**

```
                             1bp       .         .         .         .         :         .         .         .         .      100bp
Warm  LA101_cpcB_complete    MFDAFTKVVAQADARGEFLNAGQIDALSAMVAESNKRMDSVNRITSNASKIVTNAARELFDQQPALIAPGGNAYTHRRMAACLRDMEIILRYVTYAVFTG
      LA103_cpcB_complete    MFDAFTKVVAQADARGEFLNAGQIDALSAMVAESNKRMDSVNRITSNASKIVTNAARELFDQQPALIAPGGNAYTHRRMAACLRDMEIILRYVTYAVFTG
      LA117_cpcB_complete    MFDAFTKVVAQADARGEFLNAGQIDALSAMVAESNKRMDSVNRITSNASKIVTNAARELFDQQPALIAPGGNAYTHRRMAACLRDMEIILRYVTYAVFTG
      LA126_cpcB_fragment    MFDAFTKVVAQADARGEFLNAGQID---------------------------------------------------------------------------
      LA127_cpcB_complete    MFDAFTKVVAQADARGEFLNAGQIDALSAMVAESNKRMDSVNRITSNASKIVTNAARELFDQQPALIAPGGNAYTHRRMAACLRDMEIILRYVTYAVFTG

Cool  LA20_cpcB_fragment     MFDAFTKVVAQADARGEFLNAGQID---------------------------------------------------------------------------
      LA21_cpcB_fragment     MFDAFTKVVAQADARGEFLNAGQID---------------------------------------------------------------------------
      LA27_cpcB_fragment     MFDAFTKVVAQADARGEFLNAGQID---------------------------------------------------------------------------
      LA29_cpcB_fragment     MFDAFTKVVAQADARGEFLNAGQID---------------------------------------------------------------------------
      LA3_cpcB_fragment      MFDAFTKVVAQADARGEFLNAGQID---------------------------------------------------------------------------
      LA31_cpcB_fragment     MFDAFTKVVAQADARGEFLNAGQID---------------------------------------------------------------------------

                                       .         .         .         .         :         .       .172bp
Warm  LA101_cpcB_complete    DASVLEDRCLNGLRETYLALGVPGASVAEGVRKMKDAAIAIANDRNGITPGDCSALMSEIGTYFDRAAAAVA
      LA103_cpcB_complete    DASVLEDRCLNGLRETYLALGVPGASVAEGVRKMKDAAIAIANDRNGITPGDCSALMSEIGTYFDRAAAAVA
      LA117_cpcB_complete    DASVLEDRCLNGLRETYLALGVPGASVAEGVRKMKDAAIAIANDRNGITPGDCSALMSEIGTYFDRAAAAVA
      LA126_cpcB_fragment    ----------------YLALGVPGASVAEGVRKMKDAAIAIANDRNGITPGDCSALMSEIGTYFDRAAAAVA
      LA127_cpcB_complete    DASVLEDRCLNGLRETYLALGVPGASVAEGVRKMKDAAIAIANDRNGITPGDCSALMSEIGTYFDRAAAAVA

Cool  LA20_cpcB_fragment     -----------------LALGVPGASVAEGVRKMKDAAIAIANDRNGITPGDCSALMSEIGTYFDRAAAAVA
      LA21_cpcB_fragment     ----------------------------------------IANDRNGITPGDCSALMSEIGTYFDRAAAAVA
      LA27_cpcB_fragment     ----------------YLALGVPGASVAEGVRKMKDAAIAIANDRNGITPGDCSALMSEIGTYFDRAAAAVA
      LA29_cpcB_fragment     ----------------YLALGVPGASVAEGVRKMKDAAIAIANDRNGITPGDCSALMSEIGTYFDRAAAAVA
      LA3_cpcB_fragment      ----------------YLALGVPGASVAEGVRKMKDAAIAIANDRNGITPGDCSALMSEIGTYFDRAAAAVA
      LA31_cpcB_fragment     ------------------------------------------------TPGDCSALMSEIGTYFDRAAAAVA
```

Figure S6: Alignments of **A** *cpcA* and **B** *cpcB* amino acid sequences recovered from assemblies using Anvi'o. Assemblies for LA20, 27, and 29 have small fragmented terminal *cpcA* copies that are not shown here for simplicity. In each case they perfectly align with the *cpcA* copies found in the 30 °C isolates.

Supplemental Note 1:

It is possible that there could be sequence variation between these unassembled copies. For the closed genome of the warm temperature isolate LA127, variation was seen between its two copies of *cpcA* and *cpcB* (Table S5). A closer look at the read-recruitment to the complete copies of *cpcA* and *cpcB* in the draft genomes showed nucleotide positions that appear to vary between these unassembled gene copies near the 3' end (Figure S6A). This variation was in 37.8% of the reads. For *cpcB*, reads that recruit inside the gene reveal 6 single-nucleotide variant sites spread across the gene. Four of the six variant bases occur with a frequency of ~30%, and the other two at ~50% (Figure S7B).

For both *cpcA* and *cpcB* these SNVs only occur at the third base in codons and likely do not contribute to differences in peptide sequence. This is also true when looking at amino acid sequence for the copies of the *cpcA* and *cpcB* genes in the complete LA127 genome (Table S6B). Mapping reads for all isolates back to the complete genome using bowtie2's "—very-sensitive" flag did not detect any SNVs between reads and the mapped sequence for these two genes. Further, when BLASTed all completely assembled copies of both C-phycocyanin genes had a 100% match to all other complete copies in the other draft assemblies and to one of the copies in the complete genome. This suggests that although some variation exists between copies of these genes within a genome, all isolates share these differences and sequence variation for *cpcA* and *cpcB* do not explain the observed phenotypes.

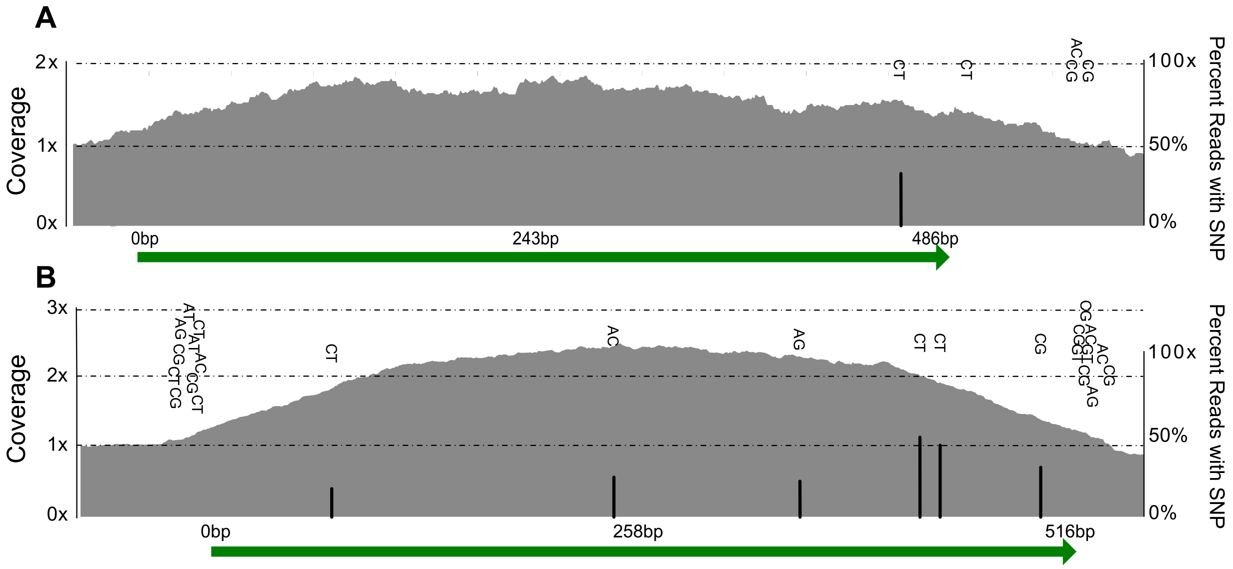


Figure S7: Visualization of reads mapped to **A** cpcA for the cool temperature isolate LA20 and **B** cpcB for the warm temperature isolate LA127. Base by base coverage is shown using grey and green arrows show the location of the genes. Single nucleotide variation between the assembled sequence and mapped reads within each gene are shown using the vertical bars and letters. Height of bars corresponds to the percent of total mapped reads that contain the SNV, indicated with the right axis.

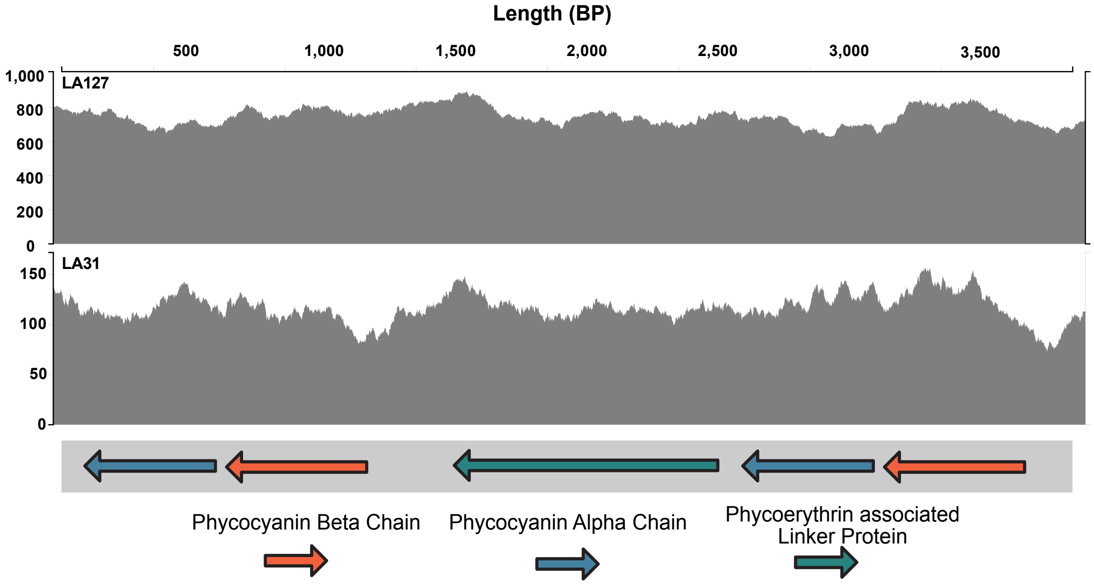


Figure S8: Results of mapping all reads to the complete genome from the warm temperature isolate LA127. Shown here is the coverage (grey shading) to the locus shown in Figure 1. The two isolates shown here are representative of all warm (top) and cool temperature isolate reads. Notably, no SNVs were shown across this locus, suggesting a high degree of sequence similarity.

Supplemental Note #2:

Looking at the *cpcA* genes that successfully assembled from the isolates derived from cool temperatures (blue arrows in Figure S8), 1 complete copy was assembled, while there are 2 copies present in the Chesapeake Bay strain CB0101 (Figure 3A) and in the hybrid assembly of the warm temperature strain LA127 (hereafter referred to as the complete genome). It is possible that variation in the copy number could contribute to distinct pattern of assembly failure. Too much similarity between gene copies can be difficult for assemblers to distinguish and could result in assemblers collapsing genes. Examining the recruitment of reads that built the assembly reveals that the single complete copy of *cpcA* has 1.6x the coverage for *cpcA* as compared to the mean coverage of the rest of the genome (Figure S6A and Figure S8B). It is likely then that all of these isolates whose draft genomes assembled a complete *cpcA* copy actually had more than one copy.

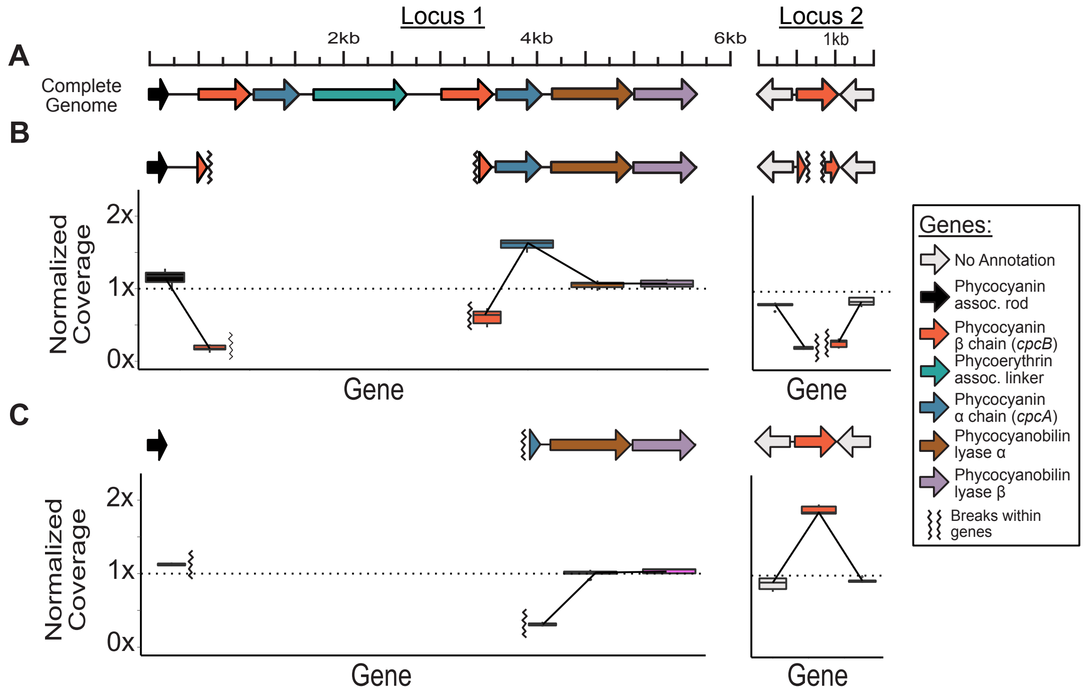


Figure S9: **A** Two loci containing genes coding for components of the accessory pigment C-phycocyanin in the closed genome for warm temperature strain LA127. **B** The same loci for all draft assemblies for isolates from cool temperatures with the results of mapping each isolate's reads back to its own assembly shown as a boxplot. The dashed line shows 1x coverage and contig breaks within genes are shown as jagged vertical lines. **C** Loci for warm temperature isolate draft genomes and coverage.

Switching focus to *cpcB* (orange arrows in Figure S8), the closed genome and closely related CB0101 genome possesses 3 copies across these 2 loci (Figure S8A). For the isolates recovered from the cooler temperatures, no copies of *cpcB* successfully assembled (Figure S8B, orange arrows). However, one *cpcB* gene did successfully assemble in four of the five warm temperature-derived isolates (Figure S8C, orange arrows). In these, read recruitment revealed mean gene- coverages of about double the mean of the rest of the genome (Figure S8C), and closer inspection of the read recruitment to this gene showed this was not uniformly distributed across it, but rather 60% of the gene had about 2.3X coverage and the rest quickly dropped to 1X approaching the 5' and 3' end of the gene. This is likely due to the variation that exists outside of the gene-copies, preventing reads from recruiting near the ends (Figure S6B). This suggests to us that while only one copy of *cpcB* assembled in these warm temperature-derived isolates, they likely possess multiple copies. Mapping reads from all isolates back to the C-phycocyanin gene copies in the closed genome did not show statistically significant differences in the relative coverage for reads from isolates of different temperatures. This suggests that they may all have similar copy numbers of these two genes; however, as another means of gaining insight into the variability of these loci, we recruited our isolate reads to the nearest relative with a closed genome, CB0101. Because CB0101 and our isolates only share 85.5% ANI (±0.04 SD), genome-wide coverage was low (12% ±0.5 SD) compared to mapping reads from each isolate to its own assembly. However, mapping rates to CB0101's copies of *cpcA* and *cpcB* were four to five times higher than to the rest of its genome, indicating these genes are more highly conserved than the rest of the genome as a whole and a good target for mapping. When mapping reads from warm and cool temperature isolates to CB0101, the combined coverage of *cpcA* genes is ~1.7x that of the coverage across each isolate's genome (Figure S9). The similarity in coverage for reads coming from both sets of isolates could suggest that both have the same *cpcA* copy number. Combined coverage of CB0101's *cpcB* gene though showed that isolates from warm temperatures (30 °C) had higher coverage relative to low temperature isolates ($p = 0.004$, Figure S9) which could suggest they have a higher copy number.

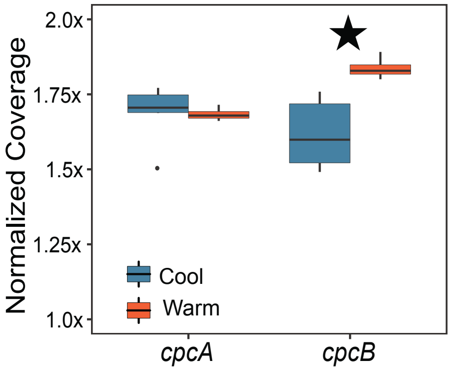


Figure S10: Normalized coverage for reads mapping to CB0101's copies of *cpcA* and *cpcB*. Star indicates significance using a student's T-test ($p < 0.5$).
